# CRISPR screen for rAAV production implicates genes associated with infection

**DOI:** 10.1101/2024.09.17.613356

**Authors:** Emily E O’Driscoll, Sakshi Arora, Jonathan F Lang, Beverly L Davidson, Ophir Shalem

## Abstract

Recombinant adeno-associated virus (rAAV) vectors are an effective and well-established tool in the growing gene therapy field, with five FDA-approved AAV-mediated gene therapies already on the market and numerous more in clinical trials. However, manufacturing rAAV vectors is an expensive, timely, and labor-intensive process that limits the commercial use of AAV-mediated gene therapies. To address this limitation, we screened producer cells for genes that could be targeted to increase rAAV yield. Specifically, we performed a CRISPR-based genome-wide knockout screen in HEK 293 cells using an antibody specific to intact AAV2 capsids coupled with flow cytometry to identify genes that modulate rAAV production. We discovered that the knockout of a group of heparan sulfate biosynthesis genes previously implicated in rAAV infectivity decreased rAAV production. Additionally, we identified several vesicular trafficking proteins for which knockout in HEK 293 cells increased rAAV yields. Our findings provide evidence that host proteins associated with viral infection may have also been co-opted for viral assembly and that the genetic makeup of viral producer cells can be manipulated to increase particle yield.

## INTRODUCTION

Recombinant adeno-associated viruses (rAAVs) are currently one of the most widely used gene delivery platforms for basic research, preclinical studies, and human gene therapies. There are currently five FDA-approved AAV-based gene therapy products (Luxturna in 2017, Zolgensma in 2019, Hemgenix in 2022, and Elevidys and Roctavian in 2023) and numerous more in clinical trials.^1^ Several aspects of rAAV vectors make them particularly advantageous for gene therapy. They can infect a broad range of cells in various tissues with defined specificities, which can be fine-tuned by the use of different serotypes.^2^ They are able to sustain stable long-term transgene expression without the risk of random and potentially oncogenic genomic alterations. Lastly, while there are immunological barriers associated with rAAV delivery,^3,4^ these vectors are derived from a widely and naturally occurring virus which is not known to cause any human diseases and is considered generally safe for clinical use, although dose-related toxicities can occur.^5,6^

Naturally occurring AAV was discovered serendipitously during lab studies of adenovirus (AdV),^7,8^ which is among the viruses that AAV can adopt for its DNA replication. AAV is a single-stranded DNA virus with a genome of ~4.7 kb packed within an icosahedral protein capsid composed of the three different subunits VP1, VP2 and VP3. The AAV genome encodes several *rep* genes required for replication, *cap* genes encoding the capsid proteins, and an assembly activating protein (AAP) flanked by inverted terminal repeats (ITRs) that promote viral replication and packaging. In the plasmid transfection platform used to produce rAAV in HEK 293 cells, all protein coding genes are removed and replaced by the delivery payload, usually gene expression cassettes with potential therapeutic values, and additional proteins involved in viral replication and assembly are expressed from separate plasmids. The resulting viral particles are replication incompetent with most of their coding capacity being utilized for transgene delivery.

One challenge for future widespread use and equitable access to AAV-based gene therapies is the cost and labor associated with rAAV production. Several of the currently approved therapies target either ocular tissue or neonates, which have helped to circumvent roadblocks associated with very high manufacturing costs. However, many of the targets that are currently being developed are aimed at adult patients using systemic administration, which will require production at scales almost intractable for widespread use. Thus, a drastic improvement of the rAAV production process is required before the promise of AAV-based gene therapies can be achieved.

It is now increasingly appreciated that rAAV particles have extensive and specific interactions with a multitude of factors in host cells. For example, efficiency of rAAV infection depends on the interactions between the capsid proteins and cell surface receptors. Particle internalization is also mediated by several host factors involved in clathrin-mediated endocytosis, cytoskeleton-mediated endosomal trafficking, endosomal escape, and multiple routes for particle degradation.^9^ Interestingly, the same factor can contribute to multiple steps in the rAAV infection life cycle. For example, AAVR, which was identified as an essential AAV receptor for multiple serotypes,^10,11^ also facilitates intracellular trafficking.^12^ As rAAV production is most often done in mammalian cells, it is likely that other host factors can enable or inhibit this process and can be fine-tuned to increase viral yield. Indeed, previous studies identified genes that when overexpressed increase production.^13,14^

Here we set out to screen for additional host factors in HEK 293 producer cells that affect rAAV production. We used CRISPR knockout screening with intracellular antibody staining specific to assembled rAAV capsids to reveal cellular pathways that negatively and positively affect production. Surprisingly, some of the top hits for which knockout decreased rAAV production were genes associated with heparan sulfate proteoglycan synthesis that were previously identified in a screen for AAV infection. We validated these results by constructing a cell line resistant to AAV infection and using it to conduct a secondary screen that identified the same top gene hits. These results suggest that AAV co-opted overlapping mechanisms for infection and intracellular viral assembly. Our screen also revealed gene knockouts that increased viral production, suggesting that these genes are involved in actively opposing viral production. These included TMED2 and TMED10, which were recently identified as organizers of large protein supercomplexes at the ER-Golgi membrane. These protein supercomplexes are responsible for the transfer of cholesterol between organelles and the remodeling of plasma membrane lipid nanodomains,^15^ suggesting that this process might repress intracellular AAV assembly. Finally, MON2 knockout also increased rAAV production and was previously implicated in HIV-1 viral production.^16^ Altogether, we identify new pathways relevant to AAV biology that can be targeted to modulate rAAV production.

## RESULTS

### Genome-wide FACS-based CRISPR screen reveals genetic modifiers of rAAV production

To enable genome-wide, FACS-based CRISPR screening for genes that impact rAAV production, we first established a workflow for quantifying AAV production using a cellular fluorescence-based readout that identified assembled capsids only. Typically, rAAV titers are measured after cell lysis and AAV particle purification, but this approach is not compatible with pooled CRISPR screening. Instead, HEK 293 cells were fixed after rAAV triple transfection and stained for flow cytometry using an antibody specific to intracellular, intact assembled AAV2 capsids (Figure 1A). To ensure that the antibody was specific to assembled AAV2 particles, mock, Rep/Cap only, and triple transfected cells were fixed, stained, and flow sorted. The Rep/Cap only condition was virtually indistinguishable from the mock control condition whereas the triple transfected condition had an appreciable percentage of fluorescent cells following optimization of staining conditions (Figure 1B, S1).

**Figure 1:**
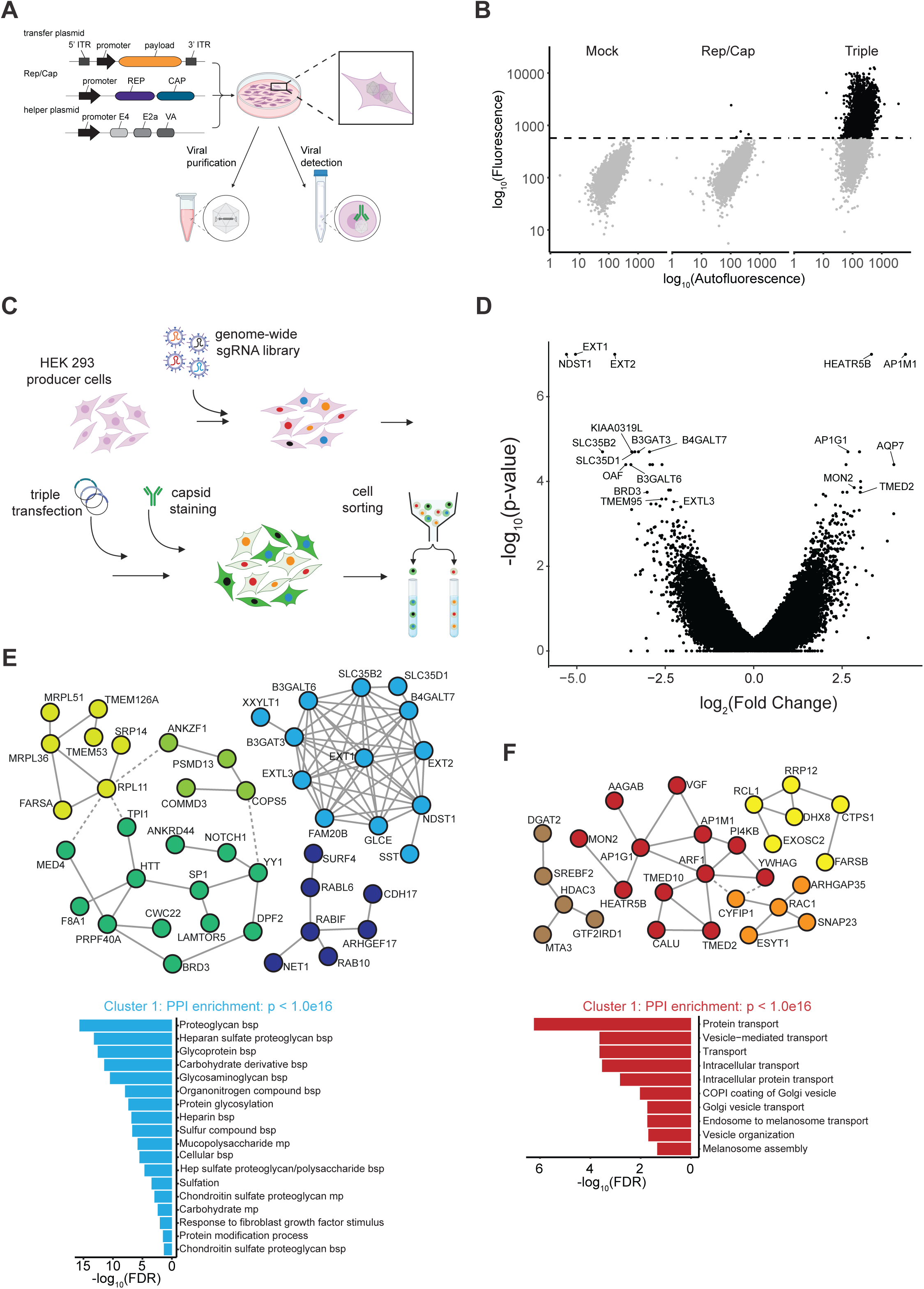
Intracellular staining for assembled rAAV2 capsids enables FACS-based measurement of rAAV production. **(A)** Schematic representation of the workflow. HEK 293 cells are transfected with a transfer plasmid, AAV2 Rep/Cap, and Ad-Helper. Following triple transfection, the cells are either lysed or fixed. Cell lysis allows for the purification and quantification of rAAV2 particles. Fixation followed by staining for intact AAV2 particles allows for detection of rAAV2 levels by flow cytometry. **(B)** Mock transfected, Rep/Cap-only transfected, and triple transfected cells were fixed and stained using a primary antibody specific to intact AAV2 capsids. Flow cytometry data validated that the antibody specifically recognized assembled capsids but not unassembled Rep/Cap. **(C)** Screening paradigm for the genome-wide FACS-based CRISPR knockout screen to identify genetic regulators of rAAV productions. HEK 293 cells were transduced first with Cas9 and then with a genome-wide sgRNA library. Following selection and expansion, the cells were triple transfected. At 72 hours post-transfection, the cells were fixed and stained for assembled AAV2. The cells were then sorted, and the least-fluorescent and most-fluorescent cells were collected for sequencing and subsequent analysis. **(D)** Volcano plot displaying the phenotype on the x axis and statistical significance on the y-axis with the top depleted and enriched genes annotated. **(E)** STRING analysis showing the protein-protein interaction clustering of the top 100 depleted genes (MCL inflation parameter = 1.5; only clusters with PPI enrichment p < 0.001 shown). Cluster 1, indicated in light blue, had significantly more interactions than expected by chance. Significantly enriched GO Biological Process terms for Cluster 1 are provided and show a strong enrichment in genes implicated in a proteoglycan biosynthetic process (bsp). **(F)** STRING analysis showing the protein-protein interaction clustering of the top 100 enriched genes (MCL inflation parameter = 1.5; only clusters with PPI enrichment p < 0.001 shown). Cluster 1, indicated in red, had significantly more interactions than expected by chance. Significantly enriched GO Cellular Component terms for Cluster 1 are provided and show an enrichment in genes involved in protein transport.

We next performed a genome-wide CRISPR-Cas9 knockout screen to identify genes in producer cells that modulate AAV production (Figure 1C). HEK 293 cells were transduced with Cas9 lentivirus and selected for five days before being transduced with a genome-wide CRISPR knockout lentiviral library^17^ at a low multiplicity of infection and selected for three days. The cells were expanded, remaining on alternating selection for a total of 10 days to ensure expression of both the Cas9 and sgRNA and to allow time for editing. Cells then underwent AAV triple transfection and were fixed and stained 72 hours later. Stained cells were subsequently sorted based on fluorescence, collecting the 15% least and most fluorescent bins. To maintain greater than 1000-fold coverage, cells were sorted over multiple days. Following genomic DNA extraction, amplicon sequencing of the sgRNA cassette was performed to measure abundance of sgRNA sequences in the two collected populations.^18^

Amplicon sgRNA read counts were analyzed to generate gene-based quantifications. The phenotypic effect was measured by calculating the fold change of the average of the two top-performing sgRNAs per gene and significance was determined by calculating a p-value using all sgRNAs per gene. The analysis revealed many genes with significant scores associated with both reduced and increased rAAV production (Figure 1D). To further investigate the pathways enriched in our primary screening results, we analyzed the top 100 gene hits in both directions using the STRING database to look for protein-protein interactions.^19^ The top cluster of genes that decreased rAAV production upon knockout was enriched in annotations related to proteoglycan biosynthesis, including several more specific terms related to heparan sulfate biosynthesis (Figure 1E). Interestingly, the majority of the highest-ranking genes for which knockout reduced rAAV production are in this cluster and are associated with heparan sulfate biosynthesis. This includes EXT1, EXT2, NDST1, B3GAT3, and others (Figure 1D-E). The top 100 gene hits that increased AAV production upon knockout were enriched in cellular component annotations related to protein transport and vesicular trafficking (Figure 1F).

### Genes involved in AAV cellular entry are also implicated in AAV production

Of the genes for which knockout decreased rAAV production, both the STRING and g:Profiler analyses^19,20^ highlighted a group of heparan sulfate biosynthesis-related genes (Figure 1E, S2). Membrane-associated heparan sulfate proteoglycans play an established role in AAV2’s ability to bind to and infect cells,^21,22^ and this same group of genes was identified in a previously published screen for genetic regulators of AAV2 cellular entry.^10^

With this in mind, we colored the depletion arm of our AAV2 production screen based on the groups of genes implicated in this previous screen for AAV2 cellular entry.^10^ Of the eight previously-implicated heparan sulfate biosynthesis genes, seven of them were among our highest-ranked gene hits. In fact, these heparan sulfate genes and the AAV receptor gene (AAVR) made up the majority of our top hits (Figure 2A). However, other genes implicated in mediating AAV2 infectivity did not appear to modulate rAAV production in our screen (Figure 2A, S3), suggesting that this result is not due to reinfection of producer cells. Still, to further exclude this possibility, we generated a clonal AAVR knockout line (Figure 2B). While AAVR is not the only gene involved in AAV2’s cellular internalization, it is required for efficient AAV2 infectivity.^10–12^ As such, transduction of our clonal AAVR knockout line with AAV containing a GFP transfer plasmid showed little green fluorescence as compared to wildtype cells (Figure 2C).

**Figure 2:**
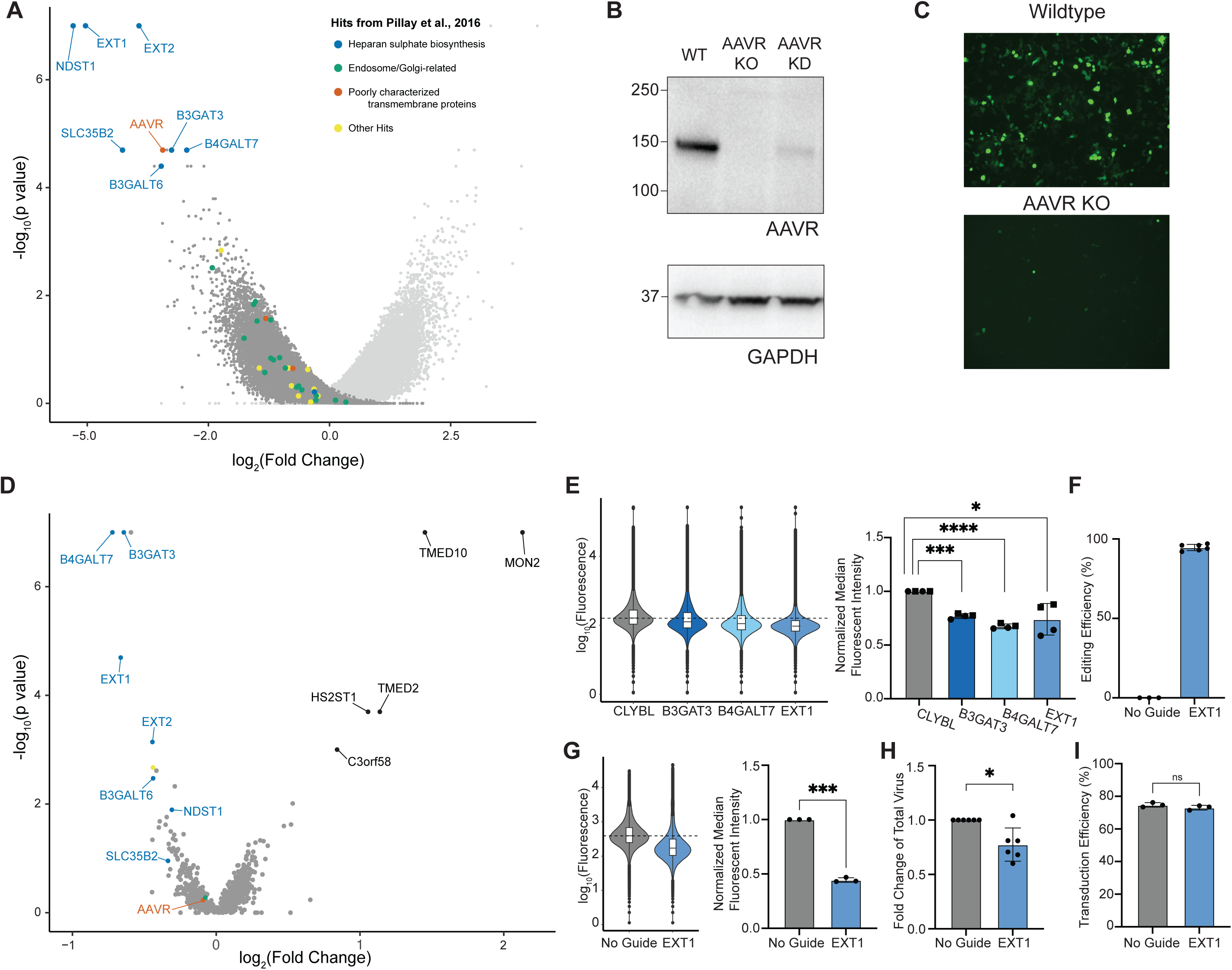
Genes implicated in AAV infection are involved in production. (**A**) The depletion arm of the rAAV production screen’s volcano plot is colored based on genes implicated in AAV infection from Pillay et al., 2016. (**B**) AAVR knockout (KO) clonal HEK 293 line confirmed by Western blot. (**C**) AAV-GFP transduction in wildtype HEK 293 and AAVR KO clonal lines. (**D**) Secondary screen of the top hits from the genome-wide rAAV production screen. The volcano plot is colored by the previously-published AAV infectivity screen. (**E**) FACS-based quantification of rAAV production in polyclonal knockout lines generated using lentiviral-delivered Cas9 and sgRNA. Two sgRNAs were tested per gene (denoted with circle and square points) in duplicate. A representative FACS plot from one replicate is shown in addition to the normalized median fluorescent intensities. (**F**) Polyclonal EXT1 knockout lines were generated by electroporation of Cas9/sgRNA RNPs and compared to HEK 293 cells electroporated with only Cas9. Editing efficiency was quantified by Sanger sequencing followed by ICE analysis. (**G,H**) rAAV production in EXT1 knockout lines as measured by FACS (G) and qPCR (H). (**I**) AAV particles produced in EXT1 polyclonal knockout lines were used to transduce wildtype HEK 293 cells and transduction efficiency was assessed by flow cytometric-based quantification.

Next, we used the clonal AAVR knockout line to perform a focused secondary CRISPR knockout screen (Figure 2D). This targeted library contained 8 sgRNAs each for the top genes identified from the genome-wide screen. The clonal AAVR knockout line was transduced with Cas9 followed by the targeted sgRNA library, and the screen was performed as described for the genome-wide screen. The only exception was that the cells were fixed 48 hours post-AAV transfection instead of at 72 hours to further reduce the possibility of reinfection.

In the depletion arm of this focused screen in AAVR knockout cells, AAVR was no longer a hit, as expected. However, heparan sulfate biosynthesis genes remained as top hits, suggesting their role in rAAV production is independent of their role in AAV2 cellular entry. We next took three of the highest-ranked heparan sulfate biosynthesis genes and tested them in an arrayed format (Figure 2E). Two sgRNAs each for B3GAT3, B4GALT7, EXT1, and the control locus CLYBL were used to knockout their respective gene in wildtype HEK 293 cells in duplicate. The lines then underwent AAV triple transfection and were subsequently fixed, processed, and flow sorted to quantity assembled AAV2 capsids levels. Compared to the CLYBL control, all three heparan sulfate biosynthesis genes tested showed a significant decrease in median fluorescence intensity, suggesting a decrease in AAV production upon knockout of either B3GAT3, B4GALT7 or EXT1.

To further validate the role of heparan sulfate biosynthesis genes in rAAV production using an orthogonal gene perturbation approach, we electroporated wildtype HEK 293 cells with RNPs containing Cas9 protein and an sgRNA against EXT1 (Figure 2F-H). As a control, we electroporated HEK 293 cells with Cas9 protein only. Our polyclonal EXT1 knockout populations showed a high level of editing efficiency (Figure 2F). AAV-transfected EXT1 knockout cells had a significant decrease in median fluorescent levels compared to control, indicating EXT1 knockout decreased AAV production (Figure 2G). Additionally, EXT1 knockout producer cells had decreased rAAV yields when quantified using qPCR (Figure 2H). To test the functionality of rAAV produced in EXT1 knockout producer cells, the viral particles produced in the EXT1 knockout lines were then used to transduce wildtype HEK 293 cells. Transduction efficiencies were indistinguishable from AAV produced in wildtype cells electroporated with Cas9 only (Figure 2I). This suggests that while knockout of EXT1 in producer cells reduces viral yield, it does not alter the functionality of the AAV particles.

### Related vesicular trafficking proteins modulate AAV production

The focused secondary screen in the clonal AAVR knockout line also brought several genes to our attention for which knockout increased rAAV yields (Figure 2D). The top ten genes consisted of known protein-protein interactors and were enriched in annotations related to vesicular protein trafficking (Figure 3A). We took the top three genes and tested them in an arrayed format. We first generated polyclonal knockout lines of TMED2, TMED10, and MON2 using lentiviral-delivered Cas9 and sgRNAs and quantified rAAV production using our FACS-based approach (Figure S4A). While some sgRNAs showed a trend towards increased rAAV production, there was large sgRNA-dependent variability likely associated with differences in gene editing efficiency and variability in small scale rAAV production.

**Figure 3:**
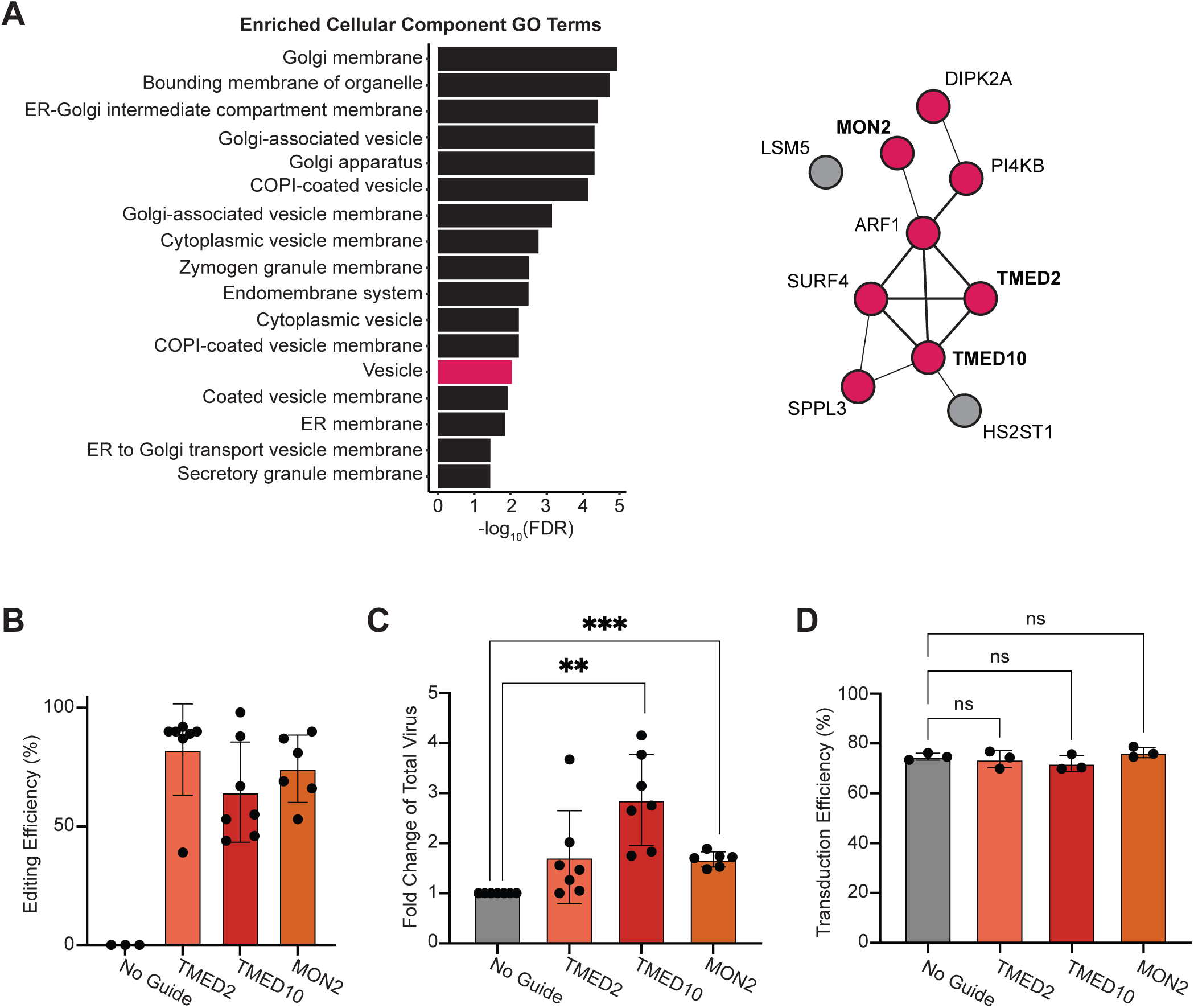
Network analysis identifies transmembrane trafficking proteins that modulate AAV production. (**A**) STRING analysis showing all potential protein-protein interactions (with interaction scores > 0.150) between the top 10 hits from the secondary screen where gene knockout increased rAAV production. Significantly enriched GO terms included many vesicle-related annotations. (**B**) Polyclonal knockout lines were generated by electroporation of Cas9/gRNA RNPs and compared to HEK 293 cells electroporated with only Cas9. Editing efficiency was quantified by Sanger sequencing followed by ICE analysis. (**C**) rAAV production as measured by qPCR in TMED10 or MON2 knockout lines. (**D**) Flow cytometric-based quantification of wildtype HEK 293 cells transduced with AAV produced in polyclonal knockout lines.

Therefore, to achieve higher KO efficiency, we tested all three genes using the orthogonal RNP-based approach described in the previous section. The editing efficiency of the TMED2, TMED10, and MON2 lines varied but averaged 82%, 64%, and 74% respectively (Figure 3B). Any lines with editing below 10% were excluded from further analysis. rAAV production in these lines was quantified using both the FACS-based and qPCR-based methods and was compared to wildtype cells electroporated with only Cas9. Knockout of either TMED2, TMED10, or MON2 trended towards an increase in fluorescence as measured by FACS (Figure S4B). When quantified by qPCR, TMED10 knockout or MON2 knockout significantly increased rAAV production (Figure 3C). Furthermore, virus produced in the knockout lines was used to transduce wildtype HEK 293 cells, and there was no significant difference in the transduction efficiency compared to when control virus was used (Figure 3D). Thus, knockout of TMED2, TMED10, or MON2 in producer cells may be a useful tool for increasing the production of functional rAAV.

## DISCUSSION

Current costs and labor associated with rAAV production are one roadblock to providing wide access to rAAV-mediated gene therapies. We hypothesized that modification of the genetic makeup of producer cells can affect rAAV yield and be used to fine-tune production. To test this hypothesis, we performed a FACS-based genome-wide CRISPR knockout screen for rAAV production using an antibody that only recognizes assembled AAV2 capsids. Interestingly, a vast majority of the genes for which knockout reduced rAAV production were associated with heparan sulfate biosynthesis and were all previously identified in a screen for rAAV infection (Fig. 2A).^10^ Other genes that were identified in the rAAV infection screen (Fig. 2A) or in other studies (Fig. S3) that were not associated with heparan sulfate biosynthesis did not appear to be associated with rAAV production, suggesting that these findings are not due to technical limitations of our experimental system. Still, to further validate the results, we generated clonal HEK 293 producer cells that lack AAVR and are thus resistant to rAAV infection. Reassuringly, conducting a secondary screen in those lines reproduced these results, suggesting that heparan sulfate biosynthesis, a process that is essential for rAAV infection, has also been co-opted by AAV for efficient intracellular assembly. More work is required to understand the mechanisms that underlie this dependency and if over activation of this pathway can be utilized for increased rAAV production.

Our screen also identified genes whose loss resulted in increased rAAV production. Indeed, several genes associated with vesicular trafficking came up as strong hits in both our primary and secondary screen. Of specific interest were TMED2 and TMED10, two genes with a well-established interaction that are components of a protein complex localized at the ER-Golgi interface and among the p24 family of proteins involved in the biogenesis of COPI and COPII-coated vesicles.^23^ Interestingly, TMED2/10 were identified in a screen for anthrax intoxication,^15^ further connecting cellular internalization pathways with intracellular assembly. Loss of TMED2/10 affected cholesterol transport between organelles and resulted in aberrant Golgi morphology. Therefore, TMED2/10 knockout may impact the ability of producer cells to degrade full or partially-assembled capsids. Future work will assess if stronger inhibition of this pathway by knockdown of more than one gene can further improve rAAV production compared to knockout of TMED2 or TMED10 alone.

One of the challenges faced in our work was that arrayed testing of rAAV production depends on many experimental factors including but not limited to cell density, growth phase, and transfection efficiency, resulting in highly variable results in small-scale arrayed experiments. In a sense, pooled screens with sufficient cell and sequencing coverage can provide more accurate ranking of gene knockouts, as cells with different perturbations grow within the same plate experiencing precisely the same culturing and treatment conditions. Because of the variability in our assay readout, validation was best achieved using RNP knockouts of TMED2/10 and MON2 compared to polyclonal lentiviral based validation studies. This approach yielded higher knockout efficiency and avoided the effect of selection on cell growth and viability.

Lastly, as our work aimed to increase rAAV yield at production scale, we envision two ways in which such data can be incorporated into standard production pipelines: The first is by engineering producer cell lines with genetically modified genomes and the second is by transient knockdowns during viral production. For the former method, since clonal selection alone can introduce large variation in cell growth and rAAV production capabilities, any derived clones with specific gene knockouts would need to be compared to a population of clones previously optimized for production without any targeted mutations. In summary, our work presents new pathways that can be co-opted to improve rAAV production, with practical applications to the adoption of AAV-gene therapies more broadly.

## MATERIALS AND METHODS

### Molecular Cloning

#### Individual sgRNAs for lentiviral-mediated CRISPR knockout

The two sgRNA sequences with the highest fold-changes in the screens were used for targeted screen validation. The forward and reverse sgRNA sequences were ordered as primers. The sgRNA oligos were phosphorylated and annealed before being inserted by Golden Gate cloning into BsmBI cloning sites in lentiGuide-Puro (Addgene 52963).

### Cell Culture

#### Maintenance

Human Embryonic Kidney 293 cells (ATCC CRL-1573) were maintained in DMEM (Gibco 11995065) with 10% FBS and 1% NEAA (Gibco 11140076). Cells were grown at 37°C with 5% CO2 to maintain physiological pH. Cells were tested regularly for mycoplasma contamination.

#### Lentiviral Generation

Human 293Ts (ATCC CRL-3216) were plated such that they would be 75% confluent at the time of transfection in plates coated with 0.1% gelatin. Between 30 to 60 minutes before transfection, the media was changed using DMEM with 10% FBS, 1% NEAA, and 1% HEPES, and 70% of the standard volume was used. Lentivirus for individual sgRNAs was prepared in 6-well plates by co-transfecting 293Ts with 1.06 ug pMDLG (Addgene #12251), 0.57 ug pMD2G (Addgene #12259), 0.4 ug pRSV-Rev (Addgene #12253), 1.06 ug plasmid to be packaged, 100 uL Opti-MEM, and 7.35 uL PEI per individual well. Lentivirus for pooled libraries was prepared in 15cm plates by co-transfecting 293Ts with 13.25 ug pMDLG, 7.2 ug pMD2G, 5 ug pRSV-Rev, 20 ug of pooled library (Brunello Library Addgene #73178 or lab-cloned secondary library), 3 mL Opti-MEM, and 136 uL PEI per plate. 5 to 6 hours post-transfection, the media was changed to DMEM with 10% FBS and 1% NEAA. 48 hours post-transfection, the supernatant was collected and filtered through a 0.45μM filter. The supernatant was aliquoted and stored at −80°C until use. Lentivirus was thawed on ice prior to transduction.

#### AAV Triple Transfection with Luciferase Assay

Human 293 cells (ATCC CRL-1573) were seeded at 40,000 cells per 12-well well one day before transfection such that they would be 80-90% confluent at the time of transfection. The next day, the cells were triple transfected with the pAd helper, pAAV2.Rep/Cap and pTransgene in 1:1:1 molar ratio (Total DNA 1.5 ug per well) using PEI Max. Briefly, the required amount of DNA was added to 10 uL (per well) of Opti-MEM. In another tube 3.0 uL of PEI Max was added to 10 uL (per well) of opti-MEM and mixed well. PEI Max/Opti-MEM was added to DNA/Opti-MEM, pipetted up and down several times, and incubated for 15 minutes at room temperature. 20 uL of transfection mix was added to each well and the cells were incubated at 37°C. 24 hours later the media of the cells was replaced with DMEM-5 (DMEM with 10% FBS and 1% Pen-strep). For the RNP experiments (Figure 2F-I and Figure 3) we included secretary Gaussia luciferase as an additional control. For this 0.1 ug of pMCSGaussia-Dura Luc Vector (Fischer Scientific) was transfected per well and 100 uL of the supernatant was collected 24 hours post transfection to perform the luciferase assay.

#### Fixing and staining AAV-transfected cells for flow cytometry

AAV-transfected cells were lifted 48 or 72 hours post-AAV triple transfection and fixed and permeabilized using the BD Cytofix/Cytoperm™ Fixation/Permeabilization Kit (554714). For the screens, 1 mL of solution/buffer was used per 10 million cells. For validation experiments, 500 uL of solution was used per well. After fixation and permeabilization according to the kit instructions, cells were incubated with primary antibody against assembled AAV2 capsids (1:50; American Research Products 03-61055), washed using BD Perm/Wash Buffer, incubated with secondary antibody (1:200; Invitrogen A-11001), washed using BD Perm/Wash Buffer, and resuspended in PBS. Cells were stored at 4C overnight before being subjected to flow cytometry.

#### Gaussia luciferase assay

Gaussia luciferase activity was measured in the supernatant according to the manufacturer’s instructions (Pierce Gaussia Luciferase Glow Assay Kit, Thermo scientific). Briefly, 20 uL of 1:20 diluted supernatant was added to a 96 well plate followed by 50 uL of the working solution. The plate was incubated at RT for 10 minutes and luminescence intensity was measured using a luminometer at a signal integration of 500ms.

#### Quantifying AAV production

At 72 hours post-transfection, cells were harvested in PBS-MK (1 mM MgCl2 and 2.5 mM KCl) buffer (1 mL per 12 well well). Cell lysis was performed using 4 freeze thaw cycles. Lysates were then treated with 50 U/ml of Benzonase for 1 hour at RT. Lysates were then centrifuged at 15,000 g for 10 minutes to remove protein and cellular debris. 2 uL of each sample was then treated with DNAse for 2 hours at 37°C followed by heat inactivation. AAV particles were lysed using 50 uL of lysis buffer and incubated at 95°C for 10 minutes. Samples were then diluted 1:500 and vector yield was calculated using qPCR using a standard curve. Primer/Probes were designed specific to the transgene.

### Cell Line Engineering

#### Polyclonal Cas9-expressing 293 cells

For the CRISPR knockout screens and all lentivirus-mediated gene knockout experiments, low-passage 293 cells were transduced with lentiCas9-Blast (Addgene# 52962). In individual wells of 6-well plates, 0.5 million 293 cells were transduced with 100 uL of lentiCas9-Blast lentivirus in media containing polybrene (Sigma TR1003G, 1:1000). 24 hours post-transduction, the transduced cells were selected using Blasticidin (5 ug/mL; Thermofisher Scientific A1113903) for 5 days.

#### Clonal AAVR knockout line

Polyclonal Cas9-expressing 293 cells were transduced with an sgRNA targeting the AAV receptor (AAVR) as described in the section above. Following puromycin selection, the cells were single-cell sorted and clones were expanded. Clonal lines were first screened using Sanger sequencing and ICE analysis to identify clones with editing at the AAVR locus.^24^ Promising clones were further screened by Western blot to identify a clone with no AAVR expression.

#### Western blot

Cells were lysed in RIPA buffer (50 mM Tris-HCl, pH 8, 150 mM NaCl, 1% IGEPAL CA-630, 0.5% sodium deoxycholate, 0.1% SDS) supplemented with 1x cOmplete Protease Inhibitors (Roche). Samples were incubated on ice for 30 minutes then spun at >18,000 x *g* for 20 min at 4°C. Supernatant protein concentration was measured using a BCA kit. Fifty micrograms were loaded into 4%–12% Criterion XT Bis-Tris gels (Bio-Rad) and transferred to PVDF membranes for blotting. Membranes were blocked for 1 hr at RT in 5% BSA in 1x TBST (137 mM NaCl, 2.7 mM KCl, 19 mM Tris Base, 0.1% Tween-20). The blot was cut into two halves at 76 kDa. The top half was incubated with mouse anti-AAVR (KIAA0319L) diluted 1:1000 in 5% BSA in TBST and lower half in mouse anti-GAPDH diluted 1:2000 in 5% BSA in TBST for 2 hours at RT followed by 3 washes. The blots were then incubated with goat anti-mouse IgG HRP (Thermo Fisher) diluted 1:10,000 in 5% BSA in TBST for 1 hour at RT. Following washes, membranes were exposed using ECL Prime Western Blotting Detection Reagent (Cytiva).

#### Validation of the Clonal AAVR knockout line

The clonal AAVR knockout HEK 293 line was transduced with AAV-GFP virus (AAV2/1-CMV-eGFP-WPRE) at an MOI of 1e5 genome copies per cell. Wildtype HEK 293 cells were transduced as a control. The cells were imaged via fluorescence microscopy at 24 and 48 hours post-transduction.

#### Clonal AAVR Knockout line stably expressing Cas9

The AAVR clonal line generated above was transduced with lentiCas9-Blast (Addgene# 52962). In individual wells of 6-well plates, 0.5 million 293 cells were transduced with 100 uL of lentiCas9-Blast lentivirus in media containing polybrene (Sigma TR1003G, 1:1000). 24 hours post-transduction, the transduced cells were selected using Blasticidin (5 ug/mL; Thermofisher Scientific A1113903) for 5 days.

#### Targeted polyclonal knockout lines generated using Cas9 and sgRNA lentiviruses

The Cas9-expressing 293 cell line was transduced with sgRNA lentivirus targeting an individual gene locus. Loci targeted included the top-ranked genes from the focused secondary screen and the CLYBL locus as a control. The two top-ranked sgRNAs from the screens were tested for each gene. Cells were transduced by adding lentivirus and polybrene (Sigma TR1003G, 1:1000) to the cells in suspension. 24 hours post-transduction, the transduced cells were selected using puromycin (1 ug/mL, ThermoFisher #A1113803) for 3 days.

#### Targeted polyclonal knockout lines generated using electroporation of RNPs

For RNP-based knockout of genes for targeted validation experiments, Cas9 and sgRNAs were ordered from IDT. Alt-R™ S.p. Cas9 Nuclease V3 was used (IDT 1081059), and the top-recommended sgRNA was chosen for each hit (IDT Alt-R® CRISPR-Cas9 sgRNA; Hs.Cas9.EXT1.1.AA, Hs.Cas9.MON2.1.AA, Hs.Cas9.TMED10.1.AA, and Hs.Cas9.TMED2.1.AA). As a control, a condition was included without an sgRNA. To assemble the RNPs, 104 pmol of Cas9, 120 pmol of sgRNA, and PBS were brought to a total volume of 5 uL. This was incubated at room temperature for 20 minutes and then put on ice until electroporating. Electroporation was then performed using the Neon transfection system (ThermoFisher Scientific). 293 cells were resuspended in Resuspension Buffer R (Neon) and then mixed with the RNP. They were then immediately electroporated with 1 pulse at 1,500 volts for 30 ms using Electrolytic Buffer E (Neon). Following recovery, cells were expanded and evaluated for genome editing and AAV production on Day 10. To assess quantitative editing, genomic DNA was isolated, and the target region was PCR amplified and Sanger sequenced. The percentage of cells edited was determined using ICE analysis.^24^ Polyclonal lines with editing efficiencies below 10 were discarded. AAV production was evaluated in the cells as described earlier.

### CRISPR Knockout Screens in 293 for AAV Production

#### Genome-wide library plasmid preparation

Brunello genome-wide sgRNA library containing an average of 4 sgRNAs per gene and 1000 non-targeting control sgRNAs was purchased from Addgene (73178). The library was transformed into electrocompetent cells (Lucigen 60242-1) and recovered at 32°C for 16-18 hours to prevent recombination. Plasmid DNA was sequenced to confirm library distribution and sgRNA representation.

#### Focused secondary library plasmid preparation

A library of 3012 sgRNA sequences was synthesized by Twist and contained 500 non-targeting sgRNAs and 2512 sgRNAs targeting the top hits from the genome-wide screen (~8 sgRNAs per gene for the top genes identified in each arm of the genome-wide screen). The pool of sgRNA sequences was PCR amplified and inserted into lentiGuide-Puro (Addgene 52963) via Golden Gate cloning. The library was transformed into electrocompetent cells (Lucigen 60242-1) and recovered at 32°C for 16-18 hours to prevent recombination. Plasmid DNA was sequenced to confirm library distribution and sgRNA representation.

#### Lentivirus titering in 293s

To ensure low MOI lentiviral transduction, library sgRNA lentivirus was titered. For the genome-wide screen, Brunello library (Addgene 73178) virus was titered in 293 stably expressing Cas9. For the focused secondary screen, the lab-cloned secondary library virus was titered in the 293 AAVR KO clonal line stably expressing Cas9. Library lentivirus was titered by plating 2×10^6^ cells per well of a 12-well plate with increasing volumes of virus mixed while the cells were in suspension along with polybrene infection reagent (Sigma TR1003G, 1:1000). Plates were spinfected by centrifugation at 1000xg for 1 hour at 37C. After approximately 16 hours, each well was split into duplicate wells: one without treatment and one treated with puromycin (1 ug/mL, ThermoFisher A1113803). After three days, cells from each well were lifted and counted, and the ratio of live cells in the +/- puromycin wells was calculated. The virus volume that achieved approximately 30% cell survival after puromycin treatment was used for the screen.

#### FACS-based CRISPR knockout screens for AAV Production

For the genome-wide screen, low-passage 293 cells expressing Cas9 were grown to ~85% confluency before being lifted and counted. To achieve >1000x coverage, 288×10^6^ cells were mixed with polybrene infection reagent (Sigma TR1003G, 1:1000) and the Brunello genome-wide sgRNA library virus at low MOI (using the titer calculated above). After thoroughly mixing, 2×10^6^ cells were plated per well in 12-well plates and spinfected by centrifugation at 1000xg for 1 hour at 37C. After overnight incubation, all of the cells were lifted, counted, and plated in 15cm plates at 7×10^6^ cells per plate. Puromycin (1 ug/mL, ThermoFisher A1113803) was added to select for transduced cells. Cells were split every 3-4 days over the next 14 days. They were maintained on alternating puromycin and blasticidin selection the entire time. At each split, cells were counted and 160×10^6^ cells were replated across 20×15cm plates. All remaining cells were discarded.

On day 14, the cells were lifted, counted, and plated to be 75% confluent 24 hours post-plating. The following day, the cells underwent AAV triple-transfection using pAAV2-Rep/Cap, pAdeno-Helper, and CBA-luciferase and PEI. 72 hours after AAV transfection, the cells were harvested, fixed, and stained for assembled AAV capsids. Cells were then filtered through a 35μm filter (Falcon, 352235) before FACS analysis and collection. Cells were gated to have a narrow range of FCS and SSC values to select for live, single cells. Autofluorescence was detected by the 405nm laser and 450/50 filter. The fluorescence of the antibody for assembled AAV capsids was detected using the 488nm laser and 515/510 filter. The top and bottom ~20% of fluorescent cells were collected. Sorted cells were pelleted and stored at −20°C until DNA extraction.

For the focused secondary screen, clonal AAVR KO 293 cells expressing Cas9 were transduced with the secondary library virus. To maintain greater than 1000x coverage, 42×10^6^ cells were mixed with polybrene infection reagent (Sigma TR1003G, 1:1000) and secondary library virus at a low MOI (as calculated in the section above). After thoroughly mixing, 2×10^6^ cells were plated per well in 12-well plates and spinfected by centrifugation at 1000xg for 1 hour at 37°C. After overnight incubation, all of the cells were lifted, counted, and plated in 15cm plates at 7×10^6^ cells per plate. Puromycin (1 ug/mL, ThermoFisher A1113803) was added to select for transduced cells. Cells were split every 3-4 days and kept on puromycin selection for the first 4 days followed by 5 days of blast selection. At each split, cells were counted and 70×10^6^ cells were replated across 7 plates. All remaining cells were discarded.

24 hours before AAV triple transfection, 7 gelatin-coated 15cm plates were plated with 15 million cells each. 45 minutes before transfection, the media was changed with 15mL using DMEM with 10% FBS, 1% NEAA, and 1% HEPES. Each plate was co-transfected with 750 uL containing Opti-MEM, 18.08 ug pAAV2-Rep/Cap, 27.68 ug pAdeno-helper, 14.23 ug CBA-luciferase, and 120 uL PEI. Media was changed 6 hours later. 48 hours post-AAV triple transfection, cells were harvested, fixed, and stained for assembled AAV capsids. Cells were sorted as described for the genome-wide screen above. A total of 6×10^6^ of the least-fluorescent cells and 7.8×10^6^ of the most-fluorescent cells were collected. Sorted cells were pelleted and stored at −20C until DNA extraction.

#### DNA Extraction, PCR amplification, and next generation sequencing

Cell pellets were thawed on ice then resuspended in 3 mL lysis buffer (50mM Tris, 50mM EDTA, 1% SDS, pH 8). After resuspension, 15 uL proteinase K (Qiagen 19131) was added to each sample. Samples were incubated at 55°C overnight. After overnight incubation, 15 uL of diluted RNase A (Qiagen 19101, 10mg/mL) was added to each sample and mixed thoroughly. Samples were then incubated for 30 minutes at 37°C. Samples were immediately placed on ice after incubation with RNaseA, where 1 mL pre-chilled 7.5M ammonium acetate was added to cooled samples to precipitate proteins. The samples were then vortexed for 30 seconds at top speed and spun at 4000xg for 10 minutes. The supernatant was then transferred to fresh tubes, where 4 mL 100% isopropanol was added to precipitate the genomic DNA. Tubes were inverted 50 times and centrifuged again at 4000g for 10 minutes. The supernatant was decanted and 3 mL 70% ethanol was added to further purify the genomic DNA. Samples were inverted ten times and spun at 4000g for 1 minute to pellet the DNA. As much of the supernatant was removed as possible before allowing the genomic DNA to air dry for 2 hours. It was then resuspended in 200 uL of nfH2O and incubated at 65°C for one hour followed by room temperature overnight to fully resuspend the DNA. DNA was then quantified by Nanodrop.

sgRNA sequences were PCR amplified with custom primers targeting the genome-integrated sgRNA backbone and containing Illumina adapters and unique barcodes for each sample to allow for multiplexing. PCR products were gel extracted and quantified by Qubit dsDNA HS assay (ThermoFisher Scientific Q32851). All samples were then pooled in equimolar ratios and sequenced using Illumina NextSeq 500/500 v2 75 cycle kit (Illumina 20024906). Amplifications were carried out with 1×8 cycles for sample index reads and 1×63 cycles for the sgRNA.

#### Screen data analysis

Raw fastq files were trimmed to remove sequences that flank the 20bp and mapped to the sgRNA library using Bowtie. sgRNA counts were then loaded to R and the following steps were performed to calculate a phenotype and p-value for each gene. Counts were first normalized by read depth by dividing read count by sample mean, multiplying by a million and adding 1 pseudocount. Next, for each sgRNA, we calculate the fold change between the least-fluorescent to most-fluorescent sample. Fold changes are corrected for increased variance at low mean values by computing a local Z score, which is calculated by ranking all the sgRNAs by mean value between the two conditions and calculating a Z score using the 2000 sgRNA window around each sgRNA. These local Z scores are then used to calculate a *phenotype* and *p-value* for each gene. *Phenotype* is calculated as the mean of the two sgRNA with the maximum absolute local Z score. *P-value* is calculated by taking the mean of all sgRNAs against a gene and comparing that to an empirical distribution of mean local Z-score generated by 100,000 permutations of gene to sgRNA associations.

## DATA AVAILABILITY STATEMENT

All data required to reproduce the results of the paper, including the full raw results from the CRISPR screens, are provided as supplementary materials.

## ACKNOWLEDGEMENTS

This work was supported by the following grants: DP2GM137416 from NIH/NIGMS, SAP#4100083086 from PA DoH and R03NS111447-01 from NINDS awarded to O.S.. Parts of Figure 1 were created with BioRender.com.

## AUTHOR CONTRIBUTIONS

Conception of this work is attributed to BLD and OS. Experiments were performed by EEO, SA, and JFL. Screen data analysis was performed by EEO. The initial manuscript draft was written by EEO and OS, and EEO, SA, BLD, and OS provided edits.

