## Supplementary material for "CRISPR screen for rAAV production implicates genes associated with infection": Supplamental

**Figure S1**

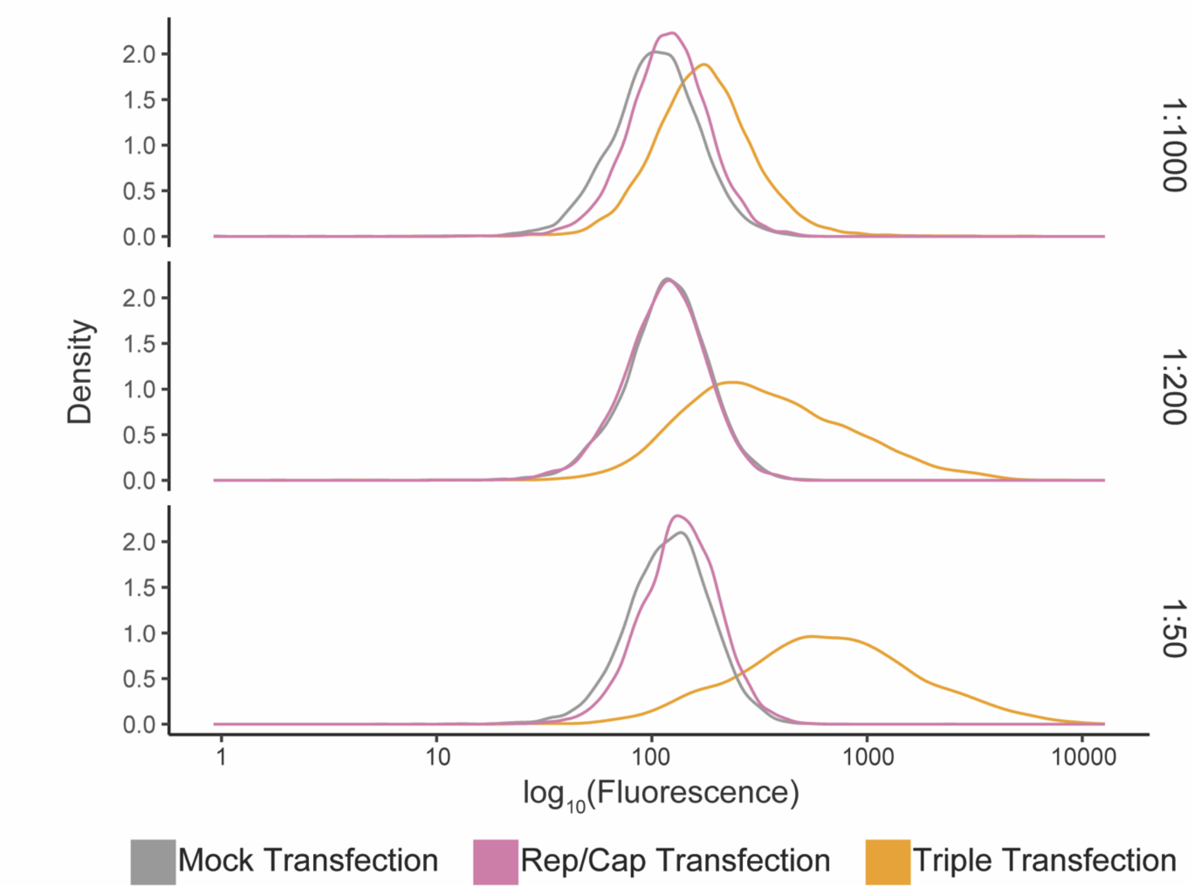


**Supplemental Figure 1: Antibody 03-61055 at a 1:50 dilution provided optimal identification of assembled AAV2 particles**. Wildtype HEK 293 cells were mock transfected, transfected with only the AAV2 Rep/Cap plasmid, or triple transfected to produce AAV. At 72 hours post-transfection, the cells were fixed and stained. A primary antibody against intact AAV2 particles (03-61055) was tested at three dilutions. All other conditions were identical between samples. Cells were then subjected to flow cytometry.

**Figure S2**

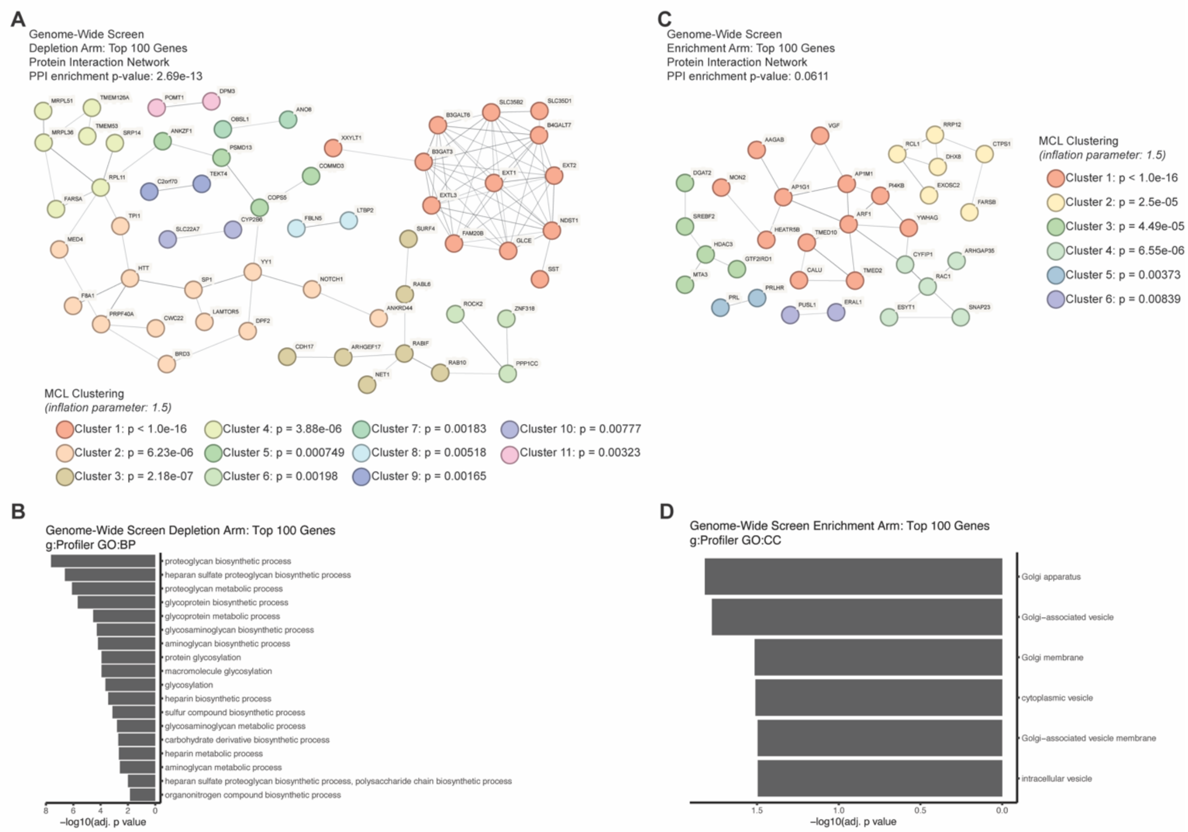


**Supplemental Figure 2: Genome-wide CRISPR screen implicates genes and pathways in modulating rAAV production.** (**A**) Complete STRING protein-protein interaction network for the top 100 genes from the genome-wide screen for which knockout decreased rAAV yields. MCL clustering was performed using an inflation parameter of 1.5. (**B**) g:Profiler analysis of the top 100 genes for which knockout decreased rAAV production revealed an enrichment of GO: Biological Process terms related to proteoglycan biosynthetic processes, particularly heparan sulfate biosynthetic processes. (**C**) Complete STRING protein-protein interaction network for the top 100 genes from the genome-wide screen for which knockout increased rAAV yields. MCL clustering was performed using an inflation parameter of 1.5. (**D**) g:Profiler analysis of the top 100 genes for which knockout decreased rAAV production revealed an enrichment of GO: Cellular Component terms related to protein trafficking.

**Figure S3**

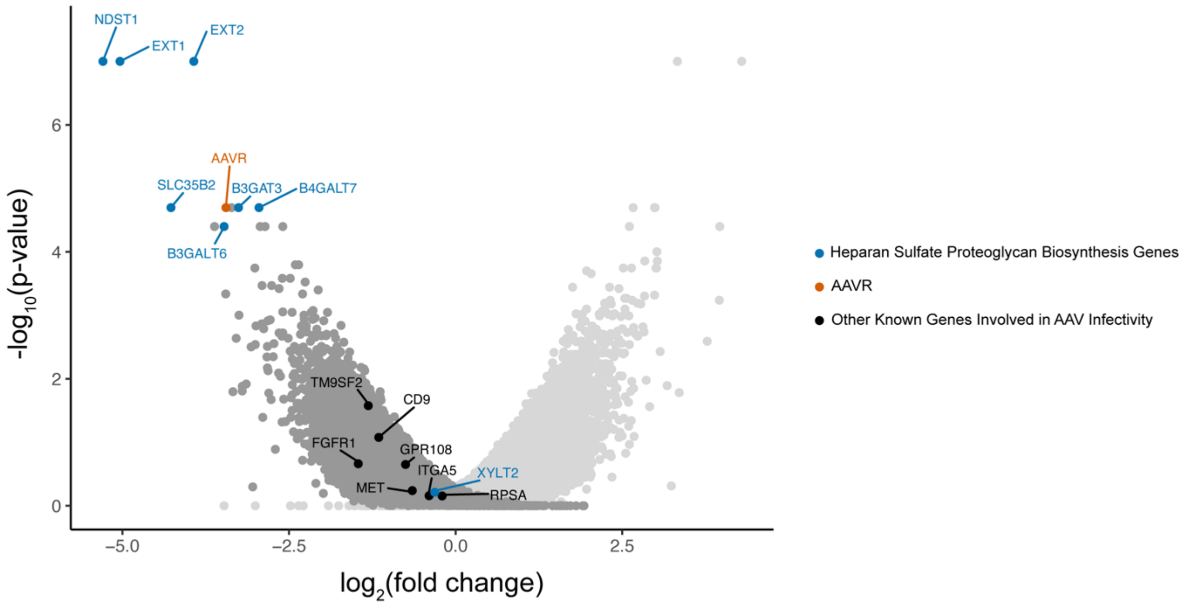


**Supplemental Figure 3: Not all genes involved in AAV2 cellular entry are hits in the rAAV production screen.** Volcano plot of the genome-wide screen for regulators of rAAV production. Among the top hits were heparan sulfate proteoglycan biosynthesis genes and AAVR, which are associated with AAV infectivity. Other genes involved in AAV2 infectivity that were not hits in our screen are labeled.

**Figure S4**

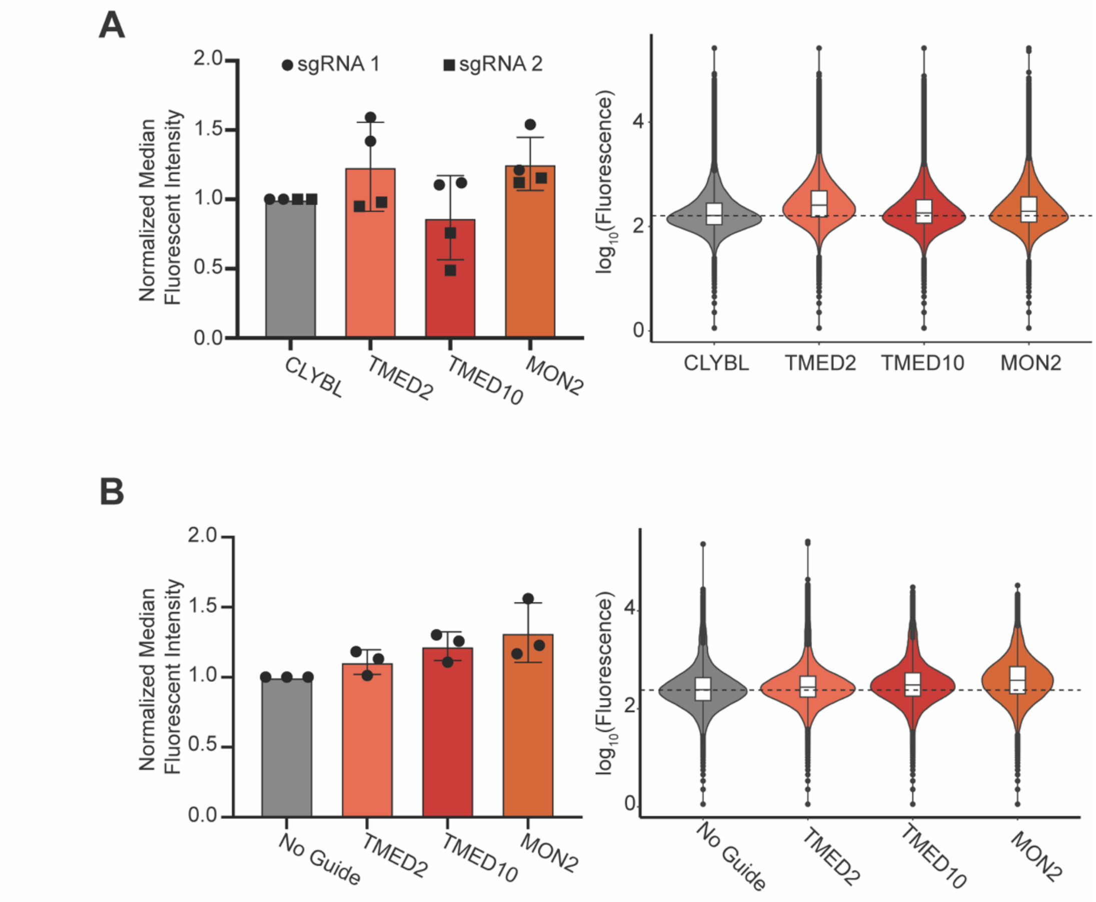


**Supplemental Figure 4: FACS-based quantifications of rAAV production in TMED10 knockout, TMED2 knockout, and MON2 knockout lines.** (**A**) Knockout lines of TMED10, TMED2, MON2, and a control CLYBL locus were generated using lentiviral-delivered Cas9 and sgRNAs. Two sgRNAs were tested per gene, and each sgRNA tested in duplicate. For both TMED2 and TMED10, there were sgRNA-dependent differences in rAAV production as measured by normalized mediant fluorescent intensity, with only one sgRNA showing a trend towards increased production. MON2 knockout showed a trend towards increased rAAV yield. A representative FACS plot using sgRNA 1 is shown. (**B**) TMED2, TMED10, or MON2 polyclonal knockout lines generated by electroporation of Cas9/gRNA RNPs trend towards increased median fluorescent intensity.

**Supplemental Table 1: Ranked Gene Hits**. The file TableS1 contains the ranked lists of all genes tested in the genome-wide and focused AAV production screens as well as their fold changes and p values.
